# Multiple stressors in the Anthropocene: Urban evolutionary history modifies sensitivity to the toxic effects of crude oil exposure in killifish

**DOI:** 10.1101/2025.02.25.640141

**Authors:** Jane Park, Charles Brown, Chelsea Hess, Madison Armstrong, Fernando Galvez, Andrew Whitehead

## Abstract

Persistence of wild species in human-altered environments is difficult, in part because challenges to fitness are complex when multiple environmental changes occur simultaneously, which is common in the Anthropocene. This complexity is difficult to conceptualize because the nature of environmental change is often highly context specific. A mechanism-guided approach may help to shape intuition and predictions about complexity; fitness challenges posed by co-occurring stressors with similar mechanisms of action may be less severe than for those with different mechanisms of action. We approach these considerations within the context of ecotoxicology because this field is built upon a rich mechanistic foundation. We hypothesized that evolved resistance to one class of common toxicants would afford resilience to the fitness impacts of another class of common toxicants that shares mechanisms of toxicity. *Fundulus* killifish populations in urban estuaries have repeatedly evolved resistance to persistent organic pollutants including PCBs. Since PCBs and some of the toxicants that constitute crude oil (e.g., high molecular weight PAHs) exert toxicity through perturbation of AHR signaling, we predicted that PCB resistant populations would also be resilient to crude oil toxicity. Common garden comparative oil exposure experiments, including killifish populations with different exposure histories, showed that most killifish populations were sensitive to fitness impacts (reproduction and development) caused by oil exposure, but that fish from the PCB-resistant population were insensitive. Population differences in toxic outcomes were not compatible with random-neutral expectations. Transcriptomics revealed that the molecular mechanisms that contributed to population variation in PAH resilience were shared with those that contribute to evolved variation in PCB resilience. We conclude that the fitness challenge posed by environmental pollutants is effectively reduced when those chemicals share mechanisms that affect fitness. Mechanistic considerations may help to scale predictions regarding the fitness challenges posed by stressors that may co-occur in human-altered environments.

## INTRODUCTION

In the Anthropocene, persistence of wild species is challenging because environmental change is often experienced severely, rapidly, and in multiple dimensions simultaneously. To persist, phenotypes (physiology, behavior, morphology) must change to suit new fitness optima. Phenotypic change may be achieved through plasticity or evolutionary adaptation. Often, the degree of human-induced environmental change is sufficient to exceed the limits of plasticity, such that fitness is impaired, and adaptive evolution becomes necessary to support persistence (Palumbi 2001, Smith and Bernatchez 2008, Hoffmann and Sgro 2011, Hendry et al. 2017). Attempts to predict the circumstances under which adaptive solutions may emerge requires engagement with the complexities (degree, pace, and dimensionality) of environmental change, including consideration of how features of environmental change interact with features of species and populations.

Features of the environment and of species interact to influence the likelihood of adaptive outcomes in human-altered environments. Relevant features of populations and species include including population size, generation time, the abundance of standing genetic variation, and the heritability and genetic architecture of relevant phenotypes (Etterson and Shaw 2001, Barrett and Schluter 2008, Bell and Gonzalez 2009, Bell 2013, Messer and Petrov 2013, Bergland et al. 2014, Kopp and Matuszewski 2014, Orr and Unckless 2014, Messer et al. 2016, Kreiner et al. 2018). Relevant features of the environment include the pace, severity, and complexity (dimensionality) of change (Tilman and Lehman 2001, Lindsey et al. 2013, Lourenço et al. 2013, Bay et al. 2017, Whitehead et al. 2017, Cisneros-Mayoral et al. 2022). Theoretical and empirical research over the past few decades have contributed much to our understanding, for example of the maximum sustainable rates of phenotypic evolution (Bürger and Lynch 1995, Hendry and Kinnison 1999, Hendry et al. 2008, Kopp and Matuszewski 2014). What is less well understood is how multiple coincident environmental changes may interact to enable or limit rapid evolutionary adaptation. A possible reason for this deficiency is that adaptation in multiple environmental dimensions may be so context-dependent that generalizable conclusions or predictions do not necessarily emerge (Orr et al. 2022). However, mechanism-guided knowledge may provide a path forward. One may predict that when co-occurring stressors share mechanisms of action, then the complexity of the fitness challenge is effectively reduced, thereby increasing the likelihood of persistence. Alternatively, stressors with different mechanisms of action could add to the complexity of the fitness challenge, and may even amplify the challenge when adaptation to one stressor is accompanied by tradeoffs in response to others (Pál et al. 2015, Faillace et al. 2021).

Pollution is a core feature of the Anthropocene (Dachs and Méjanelle 2010, Pereira et al. 2012, Dong et al. 2021), and ecotoxicology is a field with a rich mechanistic foundation. Indeed, the Adverse Outcome Pathway (AOP) framework was designed by the U.S. Environmental Protection Agency (EPA) to advance ecotoxicological risk assessment (Ankley et al. 2010). AOPs are analytical frameworks that describe chains of causality that link molecular initiating events to adverse outcomes including impacts on fitness. Ecotoxicology, therefore, provides a useful context within which to create mechanism-guided testable hypotheses about the likelihood of persistence when environmental change is complex. We hypothesize that evolved resistance to one class of environmental toxicants would provide some cross-resistance to a second class of chemicals when the two classes share features of their respective AOPs. To test this hypothesis, we included multiple populations of killifish (*Fundulus grandis*) that had different histories with, and variable sensitivity to, one class of pollutants (poly-chlorinated bi-phenyls; PCBs), and compared their sensitivity to a second class of pollutants (crude oil).

Like most fish, *Fundulus* killifish are sensitive to the toxic effects of PCBs (Nacci et al. 2010, Oziolor et al. 2014) and crude oil (Whitehead et al. 2012a, Dubansky et al. 2013, Pilcher et al. 2014, Hess et al. 2022). However, *Fundulus grandis* populations from the upper reaches of the Houston Ship Channel (TX, USA) have evolved extreme resistance to the toxic effects of persistent organic pollutants, including PCBs, following decades of extreme pollution (Oziolor et al. 2014, 2019). In vertebrate animals, PCB exposures cause multiple toxic outcomes, including cardiovascular developmental malformations during embryogenesis (Kopf and Walker 2009).

Much of the toxicity of PCBs is mediated through their activation of the aryl-hydrocarbon receptor (AHR) signaling pathway (White and Birnbaum 2009, Clark et al. 2010, King-Heiden et al. 2012, Shankar et al. 2020). Oil spills are another notorious feature of the Anthropocene (Jernelöv 2010). Crude oil is a complex mixture of thousands of chemicals, some of which are toxic, and some of which exert toxicity, including cardiovascular system developmental malformations, through their activation of the AHR signaling pathway (Billiard et al. 2008, Clark et al. 2010, Cherr et al. 2017, Incardona 2017). Because of this shared mechanism of action between PCBs and PAHs, we predicted that killifish from sites that did not have a history of exposure to persistent pollutants would be adversely affected by crude oil exposure, but that fish that had evolved resistance to PCBs would retain fitness when exposed to crude oil.

We tested whether exposures to crude oil caused fitness deficits, e.g., in adult reproduction and embryonic development, and whether sensitivity to those effects differed between populations. Since traits may vary between populations because of the influence of neutral drift or natural selection, we examined four populations where contrasts were designed to distinguish their influence. Two geographically nearby populations were sampled from within Louisiana, and two from within Texas (Figure 1). One TX population was from a relatively clean site (TX-Reference), whereas the other was from a polluted site and had evolved heritable resistance to the toxic effects of PCBs (TX-Polluted). One LA population was from a relatively clean site (LA-Reference), whereas the other was from a site that had been impacted, at least temporarily, by the Deepwater Horizon oil spill (DHOS) in 2010 (LA-Polluted). Traits evolving by neutral drift should distinguish the two TX populations from the two LA populations (e.g., because of isolation by distance, which structures genetic variation within *F. grands*; (Williams et al. 2008), see also Results). We predicted that evolved adaptive resistance to PCBs would reduce the fitness impacts of exposure to crude oil, which would be indicated by lower crude oil toxicity in fish from the TX-Polluted population compared to the others. We also tested whether the DHOS acted as an agent of natural selection by comparing sensitivity to oil exposure between the two LA populations. In addition to fitness outcomes (reproduction, embryonic development) we also measured transcriptomic responses to oil exposures in embryos to gain insight into toxicant mechanisms of action, and to reveal mechanisms that differed between populations.

**Figure 1.**
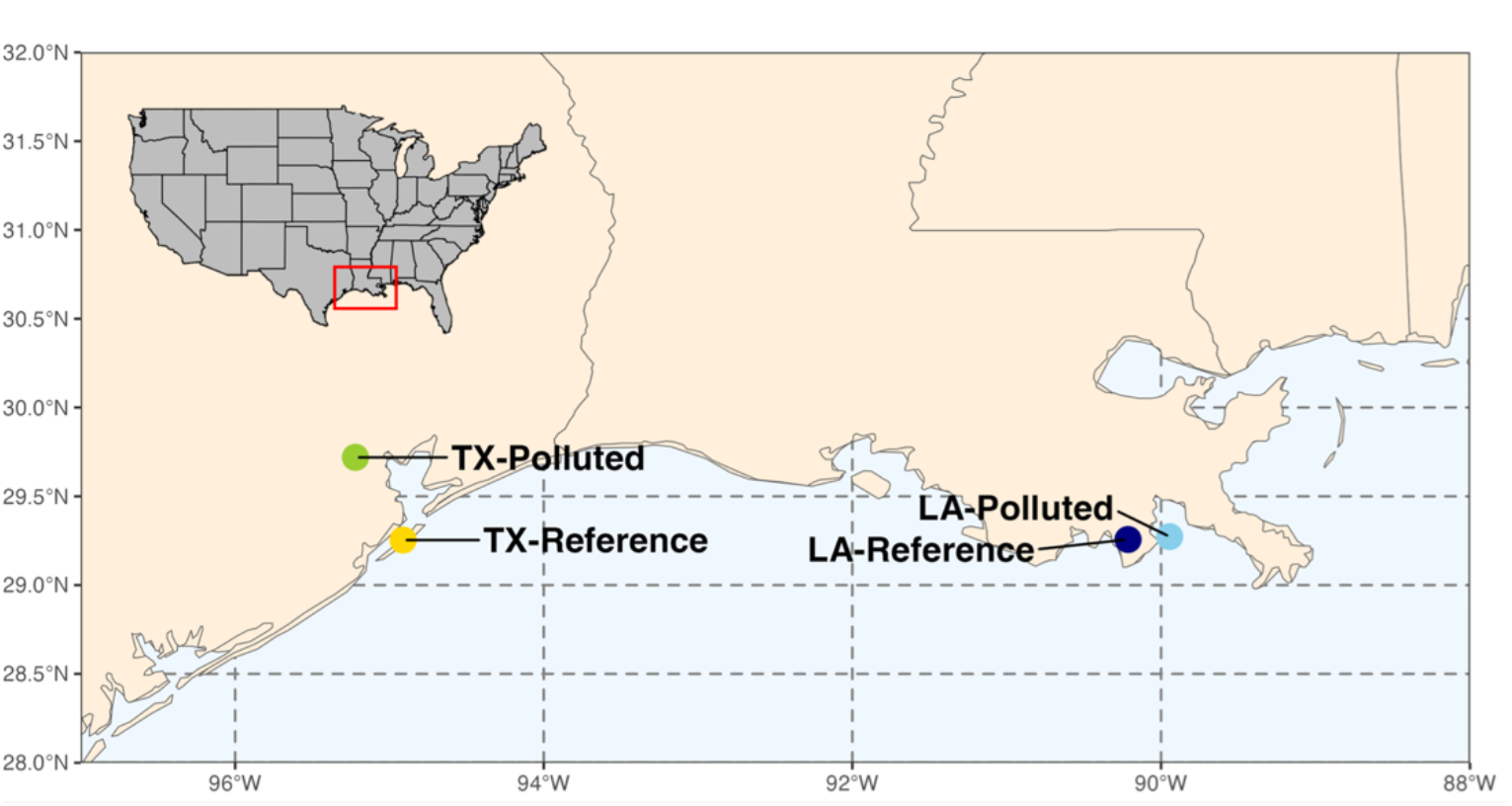
Adult *Fundulus grandis* fish were collected from four populations comprising two geographic pairs, from Louisiana (LA) and Texas (TX). Each geographic pair consists of a population from a clean reference site (−Reference) and a population from a polluted site (−Polluted). The TX-Polluted population resides in an urban estuary in the Houston Ship Channel that has been polluted with persistent organic pollutants since at least the 1970’s, where resident fish have evolved resistance to the toxic effects of PCBs. The LA-Polluted site was contaminated with crude oil during the Deepwater Horizon oil spill in 2010. To our knowledge, there is no evidence that these fish have evolved resistance to oil toxicity.

## MATERIALS and METHODS

### Adult fish collection and husbandry

All procedures that are described were performed in accordance with protocols approved by the Institutional Animal Care and Use Committee at Louisiana State University (Protocol: 15-070). Adult *Fundulus grandis* were collected from clean reference sites in Leeville, LA (LA-Reference; 29°15’24.6”N 90°12’51.3”W) and Gangs Bayou, Galveston, TX (TX-Reference; 29°15’30.31”N; 95°54’45”W; population “S1” from (Oziolor et al. 2019), and chemically polluted sites in Grand Terre Island, LA that was contaminated by the Deepwater Horizon oil spill (LA-Polluted; 29° 16’ 22.4472’’ N 89° 56’ 41.46’’ W; population “GT” from (Whitehead et al. 2012b) and (Dubansky et al. 2013)) and the polluted Superfund site at Vince Bayou, Houston, TX (TX-Polluted; 29°43’10”N; 95°13’13”W; population “R2” from (Oziolor et al. 2019)) (Figure 1). Fish from all four populations were held in common clean laboratory conditions (12 g/L artificial sea water [ASW] made with Instant Ocean salt mix; United Pet Group, Cleveland, OH, USA) at Louisiana State University for at least four months prior to experiments.

### Adult Oil Exposure

Fish from LA-Reference (n=69), LA-Polluted (n=52), TX-Reference (n=78), and TX-Polluted (n=74) were randomly assigned to control and oil treatments at an approximate ratio of 2 females to 1 male fish per replicate exposure tank. Exposures, as described below, were performed in 60-L tanks under static conditions. These adults were used to derive embryos for assessment of percent fertilization success (Figure 2). Other adults exposed to clean water alone were used as brood stock to create embryos that were used to test for oil exposure impacts on early-life development (Figure 3).

**Figure 2:**
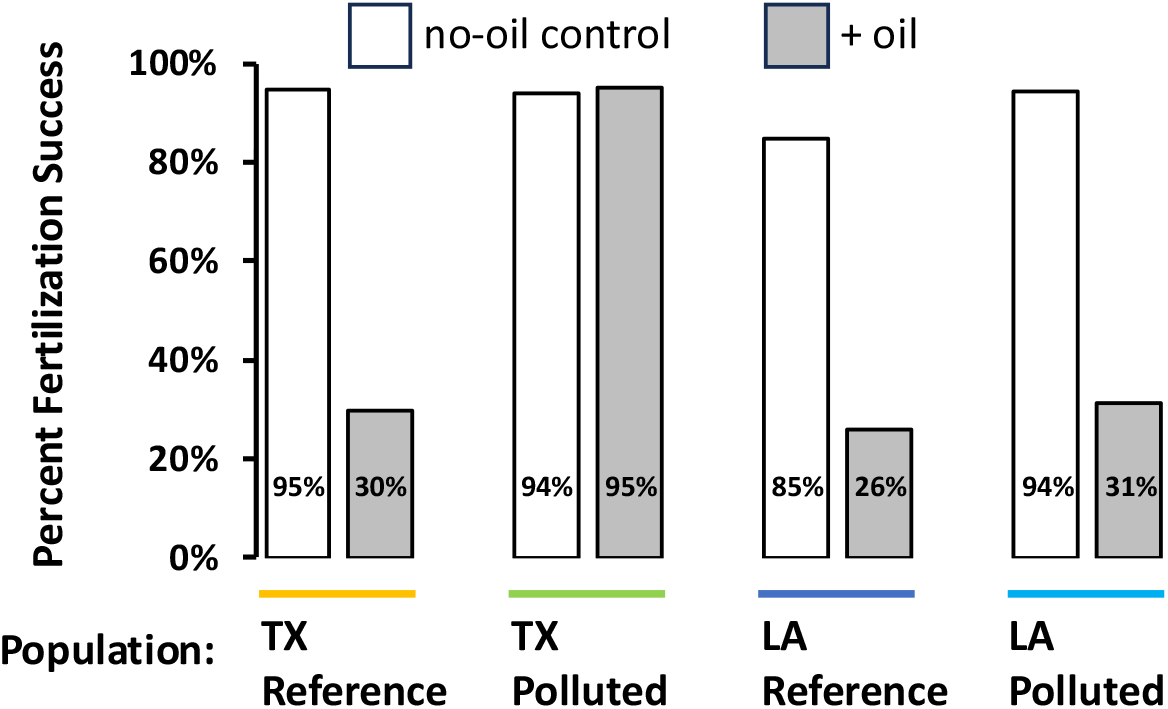
Percent of eggs with successful fertilization after adult exposure to clean water and water spiked with crude oil. White and dark filled bars indicate no-oil control or oil exposure conditions, respectively, grouped by source population. Oil exposure impaired fertilization success by nearly 70% in most populations, except for the PCB-resistant TX-Polluted population for which oil exposure did not affect fertilization success.

**Figure 3:**
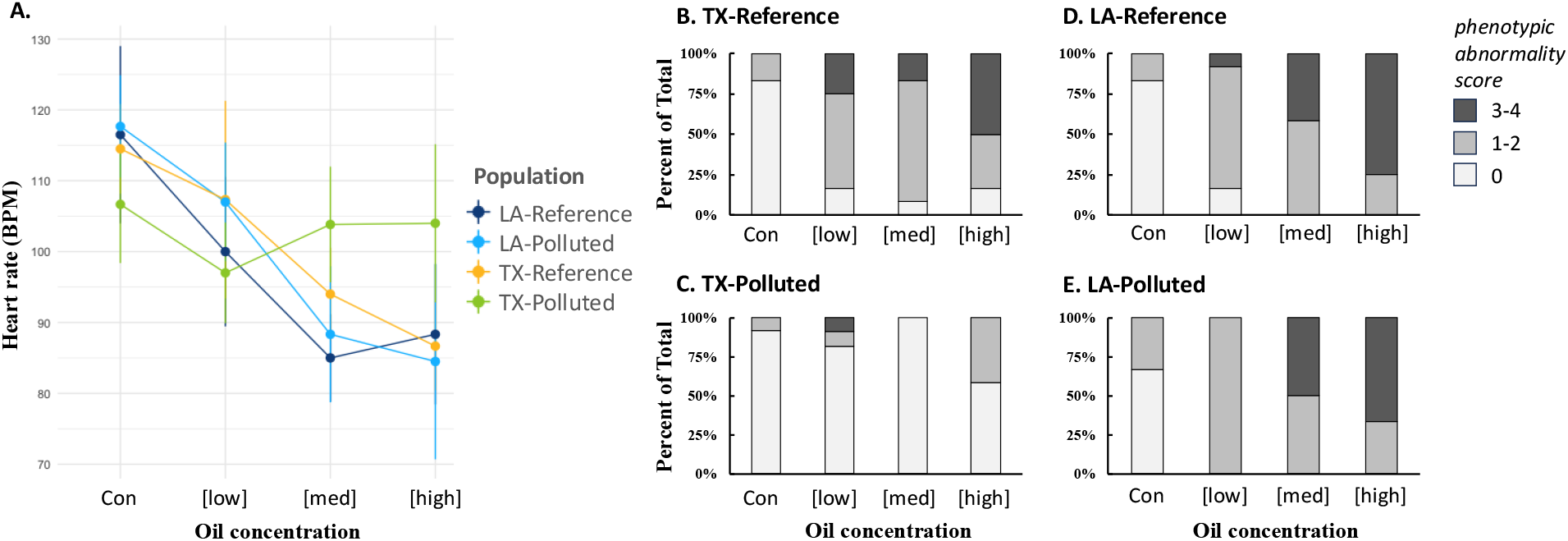
Population variation in toxicity outcomes following oil exposure during development. A: Exposure to increasing oil concentrations (x-axis, including no-oil control “con” and three increasing oil concentrations [low], [medium], and [high]) impaired heart rate (beats per minute; BPM) in a dose-responsive manner in developing embryos (7 dpf) in all populations except for the PCB-resistant TX-Polluted population for which oil exposure did not impair embryonic heart rate. B-D: Exposure to increasing oil concentrations (x-axis) induced cardiovascular system deformities in a dose-responsive manner in developing embryos (10 dpf) in all populations except for the PCB-resistant TX-Polluted population for which oil exposure did not induce deformities. Stacked bars indicate proportion of embryos within the treatment that had no observable abnormalities (PA score = 0; light gray), moderate abnormalities (PA score = 1-2; medium gray), or severe abnormalities (PA score = 3-4; dark gray).

Because of the large volume of WAF needed for adult exposures, traditional methods of producing water-accommodated fraction (WAF) of oil by high-energy mixing was not feasible. Instead, a high-volume WAF generator was constructed consisting of a 1000-liter recirculating system. Water was passed through oiled sintered glass beads (Siporax®, 15 mm, Sera) as described by Carls et al. (2000) and by Kennedy and Farrell, (2006), and the same as we reported in Park *et al*. (Park et al. 2025). The beads were pretreated with Macondo-252 surrogate light sweet crude oil (supplied by BP America Production Company; sample ID: SO-20110802-MPDF-01) at a ratio of 3.74 g oil per g bead for 48 h at 4 °C in a 2.5 L amber bottle. Oiled beads then loaded to a chamber in the upweller pipe at a ratio of 0.67 g bead L^-1^ ASW. This represented an oil loading rate of 2.5 g L^-1^ water. Beads soaked in water served as the control. Water was passed across the Siporax beads for 24 hours. The pump was turned off after 24 h allowing an oil sheen to form. The WAF was drained from the bottom to prevent the collection of surface sheen. For exposures, water from WAF tanks and clean tanks was transferred to fish exposure tanks. Every four days, ∼800 L of clean and WAF treatment waters were produced to provide 50% water exchange (30 L every 4 days) for thirty 60-L tanks (15 tanks for controls and 15 tanks for WAF exposures). Unfiltered WAF was collected from randomly selected adult exposure treatments on days 4, 20, 32, and 36 of parental exposure and control water collected on days 4 and 20. Samples were taken following 50% treatment water renewals. Adult fish from each population were exposed for 40 days to oil or no-oil control conditions.

The polycyclic aromatic compound (PAC) concentrations of water samples were analyzed as described in Park *et al*. (Park et al. 2025). Briefly, PAC analyses were performed on 150 mL composite water samples collected daily. Each composite WAF sample (*n*=6) was acidified and held at 4 °C awaiting extraction (USEPA method 3510C). Samples were shipped to ALS Environmental (Kelso, WA) for PAC analysis and alkylated homologs using GS-MS (USEPA method 8270D). A total of 6 composite samples were collected for the 40-day exposure period. The mean PAC concentration was 119.1 ± 28.3 µg L^-1^ based on the cumulative concentrations of a total of 39 PACs (see Appendix S1 for the list of 39 PACs). The analytical chemistry of water samples was used to characterize exposure conditions.

### Embryonic Oil Exposure

Embryos for developmental toxicity tests were obtained via *in vitro* fertilization from adults that had been exposed to clean water only. Eggs were stripped from adult females for each population and pooled together (separate pools for each population) in petri dishes before adding sperm from males of the same population, followed by the addition of 12 g/L ASW to activate gametes. Eggs were assessed for successful fertilization 1 hour later (hardened chorion layer and raised fertilization envelopes) and promptly moved to exposure vessels. Unfertilized eggs were removed from the study prior to initiation of exposures. Following confirmation of successful fertilization at one hour post fertilization (hpf), embryos were transferred to control or oiled water in glass dishes and reared in constant agitation for 21 days or until hatch. Water changes were performed every other day. Our experiment tested for the effects of two main factors and their interaction. The first factor was population where there were four levels (four populations), and the second factor was oil exposure where there were four levels (one no-oil control and three oil concentrations), such that there were 16 treatments total. We included four replicate exposure containers per treatment, and 12 replicate embryos per exposure container for all treatments except for embryos derived from the LA-Polluted population (only 6 replicate embryos per exposure container).

For the embryo exposures, a high energy water accommodated fraction (HEWAF) was generated from crude oil based on the protocol described in Incardona et al. (2013), and was the same as we reported in Park *et al*. (Park et al. 2025). Briefly, an initial loading concentration of 2 g of surrogate Macondo oil per liter of ASW was used to generate HEWAF preparations. HEWAF was vortexed (15,000 RPM) in a Waring CB15 blender (Waring; Torrington, CT, USA) for 30 seconds, then transferred to separatory funnels to settle. Undiluted HEWAF was then transferred to a 75-L glass aquarium and the process was repeated until a sufficient volume of WAF was produced. Undiluted HEWAF was stored at -20 °C until needed, at which point aliquots were thawed in darkness. HEWAF was then diluted to 10%, 32% or 56% with 12 g/L ASW for our three sub-lethal oil exposure conditions, equating to PAC concentrations of 53.3 ([low]), 138.0 ([medium)], and 227.9 µg/L ([high]).

### Toxicity Measurements and Analysis

For embryos, heart rates (beats per minute; BPM) of three-randomly chosen embryos per replicate container at 7 dpf were recorded by counting ventricular contractions for 30 seconds. Heart rates were analyzed using a two-way ANOVA testing for the main effects of population, oil concentration, and their interaction. Post-hoc tests (Tukey HSD) were used to identify differences between specific treatments. An additional set of embryos from each quadruplicate (*n*=3 per replicate, except where otherwise stated), were imaged 7-8 dpf with a Zeiss SteREO Lumar V.12. Brightfield images (*n*=12 per population) were taken at 36x magnification with a 0.8x objective. Images were randomly assigned numbers prior to phenotypic abnormality (PA) assessment to prevent biased evaluation. A sub-set of images (∼20% of the total number) were duplicated and their PA scores were compared to those of the originals to quantify variation in PA assessment from blind analysis (76% similarity). Embryos were evaluated for the presence of developmental abnormalities using scoring adapted from Whitehead et al. (2010), which considered abnormal morphological development, cardiac edema, heart deformation, and hemorrhaging. Embryos were assigned a 1 for each of the abnormalities when present or a 0 when absent. A cumulative phenotypic abnormality score, which ranged from 0 (none of the 4 abnormalities present) to 4 (when all 4 abnormalities were present).

### RNA-Sequencing and read count quantification

Embryos (*n* = 5 per treatment) were collected for transcriptomic analyses and flash frozen in liquid nitrogen at pre-hatch/onset of eye and pectoral fin movement (∼12 dpf) (Armstrong and Child 1965). mRNA was extracted from flash-frozen whole embryos using Zymo Directzol 96 (Cat # R2057). RNA-Seq libraries were prepared using NEBNext Directional RNA Library Prep Kit for Illumina (Cat # E7420L) which included poly-A enrichment of mRNA. Libraries were sequenced on an Illumina HiSeq 4000 as paired end 150-bp reads at the UC Davis Core Genomics Facility. Approximately 12-13 million raw reads were obtained per embryonic sample. Raw reads were checked for quality using FastQC (Andrew 2010), trimmed with trimmomatic (Bolger et al. 2014), and mapped and quantified using Salmon (Patro et al. 2017). Reads were mapped to the *Fundulus heteroclitus* reference transcriptome (Reid et al. 2017). Gene-level counts were estimated from transcript-level estimates using tximport (Soneson et al. 2016).

### Differential gene expression analysis

Genes with low read counts were excluded from RNA-Seq analysis. Low counts were defined as < 5 raw counts in < 4 replicate embryos in every treatment group, such that genes that were considered transcriptionally active (>5 counts in at least 4 replicate individuals) in at least one treatment group could be retained. We fit our data to meet the assumptions for linear regression modeling using the methods described in (Rocke et al. 2015). Log-transformed counts were normalized to the grand mean.

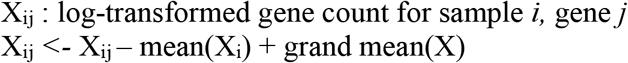

Differential gene expression was tested using a standard linear regression model in R, where we defined population and oil exposure treatment as main effects. We considered treatments effects, or their interaction, statistically significant if the FDR-adjusted p-value was < 0.05, using the following model: log(counts + *k*) ∼ Population + oil exposure + Population*oil exposure.

### Gene Ontology Enrichment Analysis

We performed gene ontology (GO) enrichment analyses on only those differentially expressed genes (DEGs) that had Uniprot entries for human or zebrafish orthologs. Clusters of DEGs with similar expression patterns (Pearson correlation) were tested for functional enrichment using DAVID Bioinformatics Resources v6.8 (updated in October 2016) (Huang et al. 2009). The background reference set included all genes that passed filtering and were included in the statistical analysis.

### SNP variant calling and population genetic summary statistics

We aligned quality-trimmed RNA-seq reads to a *F. heteroclitus* reference genome (Reid et al. 2017) (Fundulus_heteroclitus-3.0.2, RefSeq assembly accession: GCF_000826765.1) using BWA-MEM v0.7.9a (Li and Durbin 2009). We removed duplicate reads using Picard tools v2.7.1 (Broad Institute) and used Freebayes v1.3.1 (Garrison and Marth 2012) to call variants, excluding regions with low (<750) and high (>300,000) reads. We filtered variants with vcftools v0.1.16 (Danecek et al. 2011) to only include bi-allelic SNPs and quality scores > 30 (see VCF file, Appendix S2). Variants were estimated from 40, 39, 39, and 37 individuals from populations LA-Reference, LA-Polluted, TX-Reference, and TX-Polluted, respectively. We estimated genome-wide Fst values between each population pair using Weir and Cockerham’s calculations (Weir and Cockerham 1984) and vcftools v0.1.16 averaging Fst values across all variable sites.

## RESULTS and DISCUSSION

### Population genetic differentiation

To establish neutral expectations for population divergence in traits, we calculated pairwise Fst values from 2.2M variable sites sampled from 25,108 expressed transcripts. Fst between the two TX populations had previously been estimated using whole genome resequencing data (Fst = 0.024) (Oziolor et al. 2019), which is very similar to the Fst estimate from expressed transcripts reported here (Fst = 0.029, Table 1). We found lowest differentiation between populations within regions (Fst = 0.029 and 0.043 between populations within TX and LA regions, respectively), and highest differentiation between populations from different regions (average pairwise Fst across regions = 0.056). This pattern of variation within and between regions is consistent with isolation by distance and establishes the neutral context within which to consider population differences in other traits such as physiological and transcriptomic responses to crude oil challenge. For example, if populations differ in oil response traits, and those differences mirror the population genetics where divergence is smallest among populations within regions and largest between regions, then oil response trait divergence is most likely due to random-neutral processes. Patterns of trait divergence that differ from the neutral expectation may be a consequence of other evolutionary processes such as local adaptation.

### PCB-resistant fish are also resistant to oil toxicity

Adult killifish exposures to oil caused reproductive impairment, but in a population-dependent manner. Crude oil exposures caused a nearly 70% decrease in fertilization success in all populations except for the PCB-resistant TX-Polluted population, for which exposures did not perturb fertilization success (p<0.001, Fisher’s exact test, Figure 2). For this population, fertilization success remained high in both control and oil exposed treatments (94% and 95%, respectively). This pattern of population variation is not consistent with random-neutral drift but is consistent with the hypothesis that evolved PCB resistance provides cross-resistance to the reproductive impacts normally caused by crude oil exposure.

Exposures to crude oil during early-life development were not lethal but did cause perturbations of cardiovascular development and heart function in embryos. These outcomes were population-dependent. Increasing concentrations of crude oil caused a dose-dependent decrease in heart rate in all populations except the PCB-resistant TX-Polluted population (significant exposure by population interaction, p<0.001, Figure 3A). Exposures also induced dose-dependent deformities in the developing cardiovascular system in embryos from all populations except the PCB-resistant TX-Polluted population (significant exposure by population interaction, p<0.01, Figure 3B-3E). The PCB-resistant TX-Polluted population was insensitive to the effects of crude oil exposure on heart rate (Figure 3A) and cardiovascular system deformities (Figure 3C). This pattern of population variation is not consistent with random-neutral drift but is consistent with the hypothesis that evolved PCB resistance provides cross-resistance to the developmental impacts normally caused by crude oil exposure.

### Population variation in transcriptomic response to oil exposure rejects the neutral expectation

After filtration, we conducted differential gene expression analysis for 25,018 genes (Appendix S3). We detected 8,032 genes that were differentially expressed between populations (Figure S1). Excluding genes that also varied with oil exposure or had significant exposure by population interactions (p > 0.1 for oil treatment effect or interaction effect) left 5,167 genes that varied *only* among populations (e.g., no other treatment effects). The pattern of population variation for these genes is consistent with our neutral expectations, where population variation is minimal within geographic regions, but greatest between regions (Figure 4).

**Figure 4:**
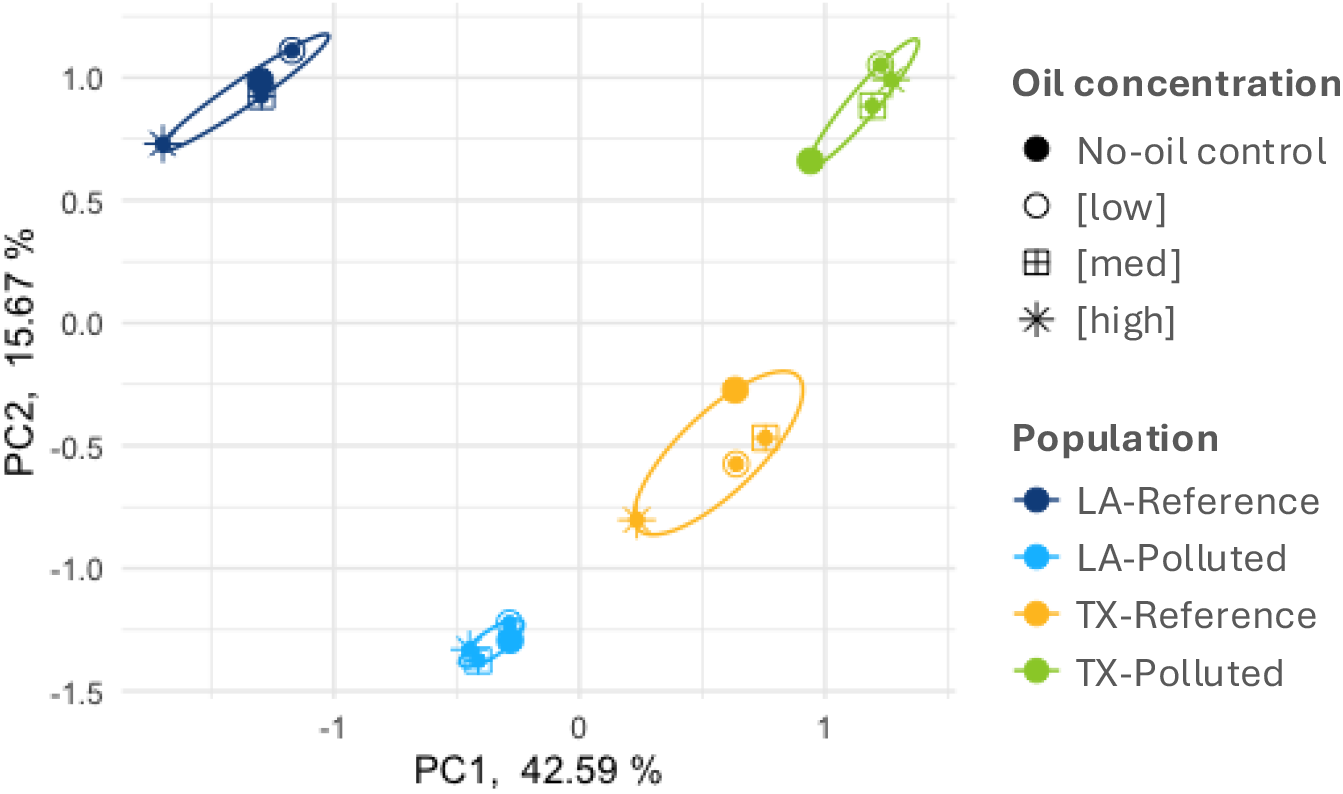
Principal components analysis (PCA) for genes that varied in expression between populations. We included 5,167 genes that varied in expression between populations, which excluded genes that also varied with oil exposure or had significant exposure-by-population interactions (p < 0.1). PCA was performed using mean expression values among replicate individuals (n=5) within each treatment (oil concentration by population).

Genes that were transcriptionally responsive to oil showed a pattern of population variation that rejected random-neutral expectations. Hundreds of genes had a transcriptional response to oil that varied between populations (512 genes with significant exposure-by-population interaction, p<0.05). For these 512 genes, the predominant pattern of population variation was a conserved response to oil exposure between the two regional reference populations (TX-Reference and LA-Reference; Figure 5). Compared to the reference populations, the PCB-resistant TX-Polluted population showed the most divergent oil exposure response. The LA-Polluted population diverged more subtly from the two reference populations, mainly at the highest oil concentrations. This pattern of population variation is not consistent with random-neutral drift but is consistent with the hypothesis that transcriptional responses to oil exposure are related to the resistance to oil toxicity that is afforded by evolved PCB resistance in the TX-Polluted population. Furthermore, since exposure-induced expression in the DHOS-exposed LA-Polluted population differs subtly from the two regional reference populations, this may indicate subtle evolved divergence in oil-responsiveness in this population.

**Figure 5:**
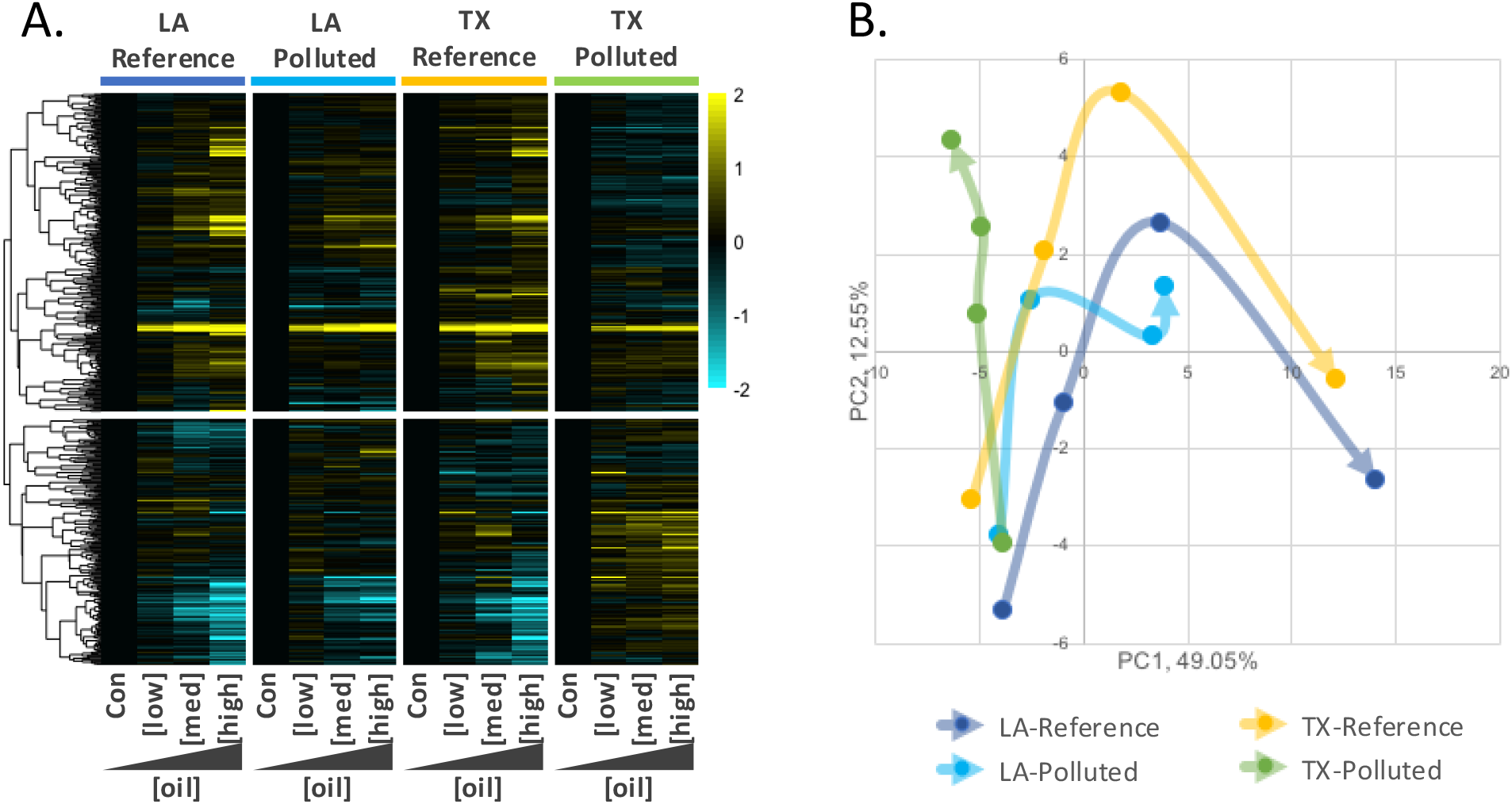
Population-dependent transcriptional responses to embryonic oil exposure. A) Heatmap includes genes that had oil exposure effects on transcription, but where those effects varied between populations (512 genes with significant oil exposure-by-population interaction, p<0.05). The four panels, left to right, show oil concentration-responsive genes for the LA-Reference, LA-Polluted, TX-Reference, and Tx-Polluted populations, respectively. Within each population panel, increasing oil concentrations are organized starting from no-oil controls (Con) on the left to the highest concentration of oil on the right. Individual genes are the rows. Genes (rows) were hierarchically clustered (Pearson correlation). For each gene within each population, expression was normalized to the control condition (black), where up-regulation and down-regulation relative to control conditions is indicated in yellow and blue, respectively. Color intensity relates to fold-increase or decrease of log2 expression (see color scale bar). B) The first two principal components for the 512 genes with significant oil exposure-by-population interaction (p<0.05). Arrows indicate the trajectory of gene expression change with increasing dose for each population. The base of each arrow represents the no-oil control condition for that population, where the arrow trajectory tracks gene expression change with increasing oil concentrations, such that the tip of arrow represents the highest oil concentration. Populations share similar gene expression under control conditions, but expression diverges between populations as the concentration of oil exposure increases. The PCB-resistant TX-Polluted population shows the most divergent response to oil. The DHOS-exposed LA-Polluted population shows a blunted response to oil at the highest oil concentrations compared to the two regional reference populations that showed nearly identical transcriptional responses to oil exposure.

### Molecular mechanisms associated with resistance to oil toxicity are shared with mechanisms that underlie evolved PCB resistance

We performed GO enrichment analysis for the 512 genes showing significant oil-by-population interaction to explore potential mechanisms underlying cross-tolerance to oil exposure (Fig. S2). We observed an enrichment of cytochrome p450-related genes, which is indicative of altered AHR signaling, which plays a key role in derived pollution adaptation in killifish (Meyer 2002, Nacci et al. 2010, 2016, Oziolor et al. 2014, 2019, Reid et al. 2016, Whitehead et al. 2017, Miller et al. 2024). Functions related to the vascular endothelial growth factor receptor signaling pathway and EFG-like calcium binding were enriched in these DEGs. Genes relating to these functions may play a role in resistance to cardiotoxicity (Zhao et al. 2010). Additionally, we observed an enrichment of neurogenesis and neurodegeneration pathways. Previous studies have similarly found impaired neural development in zebrafish (de Soysa et al. 2012), red drum (Xu et al. 2017), and mahi-mahi (Xu et al. 2016), as a consequence of exposure to crude oil.

In addition to those biological functions suggested by GO analysis above, we more closely inspected responses of the AHR signaling pathway since is widely known for its role in DLC toxicity for developing vertebrate animals (Denison and Nagy 2003, Mandal 2005, Billiard et al. 2006, King-Heiden et al. 2012). Dioxins, DLCs, and many PAHs are ligands that bind to AHR protein in the cytoplasm of cells. Ligand-bound AHR then translocates to the nucleus, forms a heterodimer with ARNT, and activates transcription of a battery of genes (Beischlag et al. 2008, Fader and Zacharewski 2017). Blockage of this AHR response protects from DLC and PAH toxicity (Clark et al. 2010). Genetic and transcriptomic variation in the AHR pathway is consistently associated with adaptive resistance to DLC and PAH pollution in multiple populations and species of killifish (Whitehead et al. 2012b, Reid et al. 2016, Oziolor et al. 2019, Miller et al. 2024) and Atlantic tomcod (Wirgin and Waldman 2004), including for the TX-Polluted population of *F. grandis* (Oziolor et al. 2019). We tested whether the AHR signaling pathway is similarly implicated in conferring cross-tolerance to crude oil in the TX-Polluted population. We identified 35 genes that are components of the AHR signaling pathway. Most of these genes (29 of 35) showed significant variation in gene expression between treatments (statistically significant main effects of oil exposure, population, or their interaction). Visualization of expression patterns for these 35 genes show a common pattern of transcriptional response to oil exposure in all populations except the TX-Polluted population (Fig. 6a, b). The expression patterns we observed for these genes, such as AHR1a, AHR2a, AHRR, and several cytochrome P450 genes, are consistent with transcriptional responses of AHR pathway genes following DLC exposure that differ between DLC-resistant and sensitive populations of killifish (Whitehead et al. 2010, 2012b, Reid et al. 2016, Oziolor et al. 2019). We conclude that desensitization of the AHR response that is adaptive in DLC-resistant populations also contributes to resistance to crude oil toxicity.

**Figure 6:**
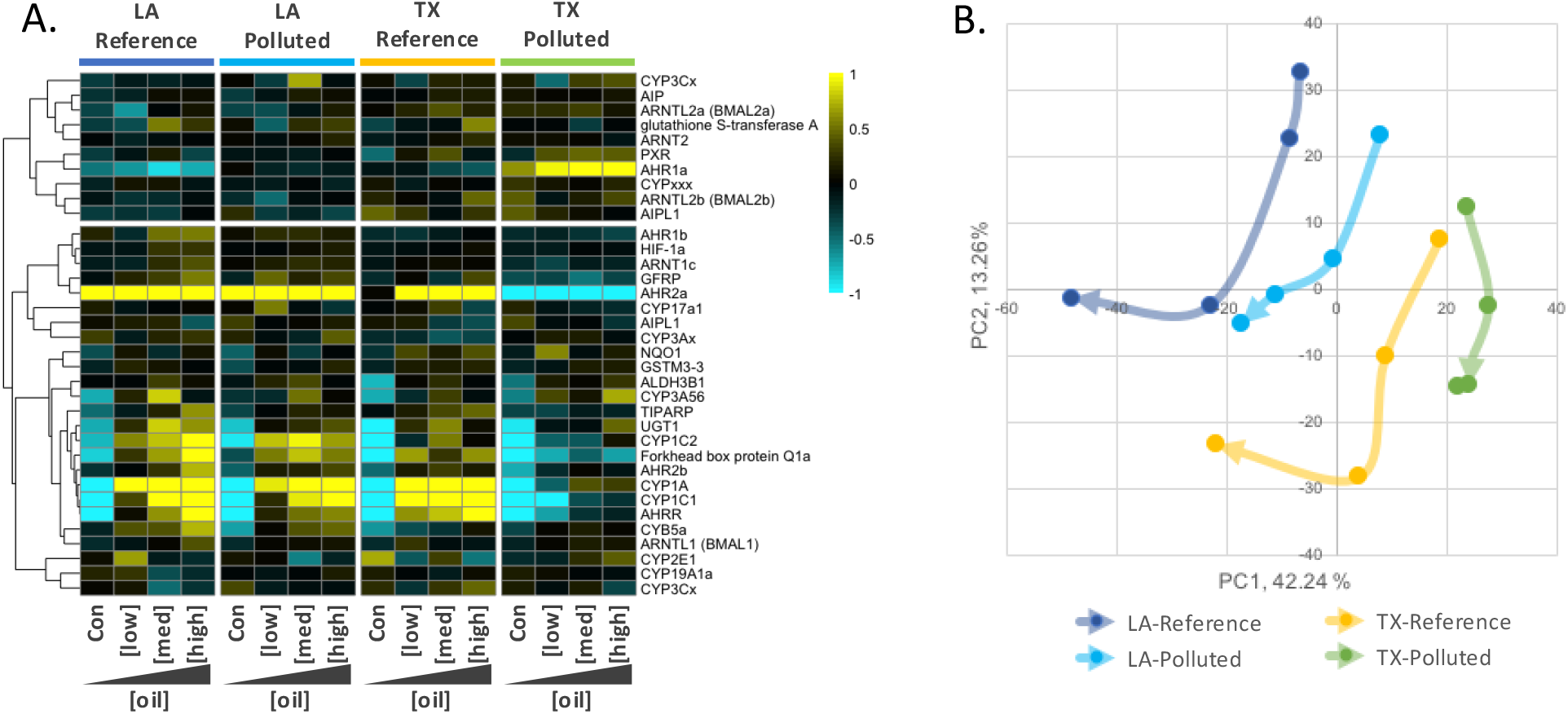
Population variation in the transcriptional responses to oil exposure for genes involved in the AHR signaling pathway, which is a key pathway that mediates toxicity. A) Heatmap includes genes that are components of the AHR signaling pathway. The four panels, left to right, show oil concentration-responsive genes for the LA-Reference, LA-Polluted, TX-Reference, and TX-Polluted populations, respectively. Within each population panel, increasing oil concentrations are organized starting from no-oil controls (Con) on the left to the highest concentration of oil on the right. Individual genes are the rows. Genes (rows) were hierarchically clustered (Pearson correlation). Higher and lower transcript abundance is indicated in yellow and blue, respectively. Color intensity relates to fold-increase or decrease of log2 expression (see color scale bar). B) The first two principal components for the AHR-regulated genes. Arrows indicate the trajectory of gene expression change with increasing dose for each population. The base of each arrow represents the no-oil control condition for that population, where the arrow trajectory tracks gene expression change with increasing oil concentrations, such that the tip of arrow represents the highest oil concentration. Populations tend to share similar AHR-mediated gene expression responses to oil exposure, except for the PCB-resistant TX-Polluted population which shows a divergent response to oil. The LA-Polluted population also shows a blunted response especially at higher oil concentrations.

We sought to identify genes with oil-responsive expression that was conserved across all four populations. We found 4,109 DEGs that show a consistent transcriptional response to oil in all populations. We did this by selecting genes with significant oil exposure-response (adj. p < 0.05) but with no significant dose by population interaction. However, p > 0.05 for a treatment effect does not equate with no treatment effect. In fact, there is no clear statistical test for “no effect”. Therefore, in order to bias our results towards genes with conserved responses across populations we chose a threshold of p > 0.1 to exclude genes with an oil dose-by-population interaction. Qualitatively, this set of genes show oil dose-dependent patterns of expression that are mostly shared across populations, though the dose response still tends to be diminished in the TX-Polluted population compared to the others (Fig S3a, b). Therefore, some of these genes may be involved in the resistant response observed in TX-Polluted site fish. However, this set of genes is enriched for those that show an oil response that is conserved regardless of whether the population is sensitive or resistant to oil-induced developmental cardiac toxicity. Gene ontology (GO) enrichment analysis (Fig. S3c) implicates oil-responsive genes involved in cardiac function, visual perception, Wnt signaling, and voltage-gated calcium ion channel function; these functions have been associated with some aspects of crude oil toxicity in killifish and many other species (Garcia et al. 2012, Fairbairn et al. 2012, Pilcher et al. 2014, Sørhus et al. 2016).

There is a substantial transcriptomic response to oil exposure in the TX-Polluted population despite tolerance to developmental cardiac toxicity (Fig. S3). We hypothesize that while these DEGs are transcriptionally responsive to the experience of oil exposure, they are not involved in the specific mechanism of cross-tolerance that protects fish from the TX-Polluted population from developmental cardiotoxicity. In contrast, these DEGs are likely part of a more universal defensive or compensatory response to exposure. These DEGs could also indicate oil responses in the TX-Polluted population that underlie phenotypes that are equally as sensitive to oiling as the other populations; although we tested for variation in developmental cardiotoxic endpoints (Fig. 3), because that is one of the most sensitive toxic responses to DLCs and crude oil in vertebrate animals (White and Birnbaum 2009), crude oil also causes toxicity through perturbations of other phenotypes (Grosell and Pasparakis 2020). It is therefore plausible that TX-Polluted fish are equally sensitive as other populations to other aspects of oil toxicity. Furthermore, the perturbations in gene expression observed here may not manifest as phenotypic perturbations until later in life. Future studies may explore oil responsive phenotypes beyond cardiac form and function during embryogenesis, such as other aspects of cardiac performance, skeletal malformations, cholesterol biosynthesis, nervous system function, or ion regulation, especially in larvae or adult fish.

We do not find clear evidence that exposure to the DHOS event caused adaptive changes in the population recently exposed to the DHOS (LA-Polluted), insofar as fish from the LA-Polluted site are as sensitive as reference populations to oil exposure-induced reproductive and developmental toxicity (Fig. 2, 3). The LA-Polluted population presumably shares many biological attributes (e.g. genetic diversity, generation time, genetic architecture of traits impacted by contaminant exposure) with fish from the TX-Polluted population, but it is possible that the genetic variants that enabled rapid adaptation in the TX-Polluted population are absent in the LA-Polluted population. Differences between the environmental challenges faced between these populations may also explain, in part, the persistent sensitivity to oil exposure in LA-Polluted site fish. It is possible that the selection pressure imposed by the DHOS was not adequately severe, lasting, or temporally consistent to cause detectable evolutionary change. Consistent with this, genome scans did not identify selective sweeps in DHOS-exposed populations of *F. grandis* (Schaefer et al. 2018). In contrast, the Superfund site in the Houston Ship Channel occupied by the TX-Polluted population has been persistently contaminated with toxic levels of DLCs since at least the 1970’s (Yeager et al. 2007). Although the response to oil exposure in LA-Polluted site fish was not as divergent as fish from the TX-Polluted site, qualitative assessments of gene expression patterns for genes with a significant oil exposure-by-population interaction (Fig. 5a, b), and AHR pathway-related genes (Fig. 6a, b), are suggestive of dampened sensitivity to higher concentrations of oil in the LA-Polluted population compared to the reference populations. This could indicate subtle evolved changes that reflect a shift toward an adaptive response in the LA-Polluted population, and merits further investigation. Whole genome population genomics data coupled with sensitive methods for detecting subtle signatures of polygenic natural selection (e.g., (Buffalo and Coop 2019, 2020)) in DHOS-exposed populations could provide useful insights. It is also plausible that DHOS exposure induced non-genetic mechanisms of trans-generational inheritance that blunted that population’s response to additional oil exposure. However, our previous experiments show that oil exposure in *F. grandis* tends to have trans-generational impacts that are maladaptive and that do not affect the molecular response to direct oil exposure (Park et al. 2025).

We adopted a mechanism-guided approach to generate and test the hypothesis that evolved resistance to multiple unrelated agents of environmental change is likely when there exists mechanistic overlap between the agent’s mode of action. Our results support this hypothesis in killifish, where evolved resistance to industrial pollution afforded by adaptive desensitization of AHR signaling provides fitness advantages when challenged with oil spills that also exert toxicity through AHR signaling. Other studies have considered a mechanism-guided approach to predict cross-resistance on the one hand, and tradeoffs (e.g., from negative pleiotropy) on the other. For example, previous research tested the prediction that adaptive desensitization of AHR signaling in killifish would confer cross-resistance to pesticides that require metabolic activation of toxicity through activation of AHR signaling, and some results were consistent with this prediction, but others were not likely because of incomplete knowledge of the complexity and functional redundancy of metabolic pathways (e.g., (Clark and Di Giulio 2012, Oziolor et al. 2016)). Similarly, because resistance to other stressors requires intact AHR signaling, other studies have predicted costs and tradeoffs associated with evolved AHR desensitization, and some results have been consistent with this (e.g., (Meyer and Di Giulio 2003, Harbeitner et al. 2013)). The pesticide and antibiotic resistance literature abounds with examples of evolved cross-resistance among selective agents (poisons) with shared mechanisms of action (Soderlund and Bloomquist 1990, Périchon et al. 2019, Colclough et al. 2019, Bras et al. 2022). A similar mechanism-guided approach applied to other species struggling their way through the multifarious fitness challenges imposed by the Anthropocene may be helpful for predicting the likelihood and pace of adaptive outcomes.

## ACKNOWLEDGEMENTS

Jennifer Roach assisted with transcriptomics data collection. Dr. Cole Matson shared fish from Texas populations. Dr. Christopher Green help with fish housing and rearing.

## SUPPLEMENTAL FIGURES

**Figure S1.**
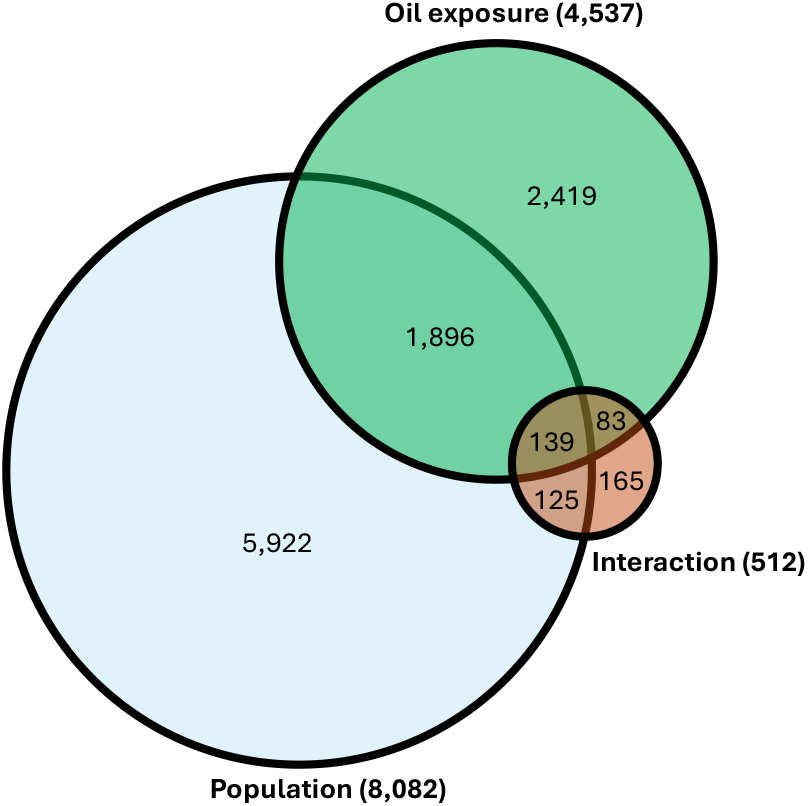
Numbers of differentially expressed genes (FDR adj. p < 0.05) for the main effects of *population* and *oil exposure concentration* treatment groups, and their interaction.

**Figure S2.**
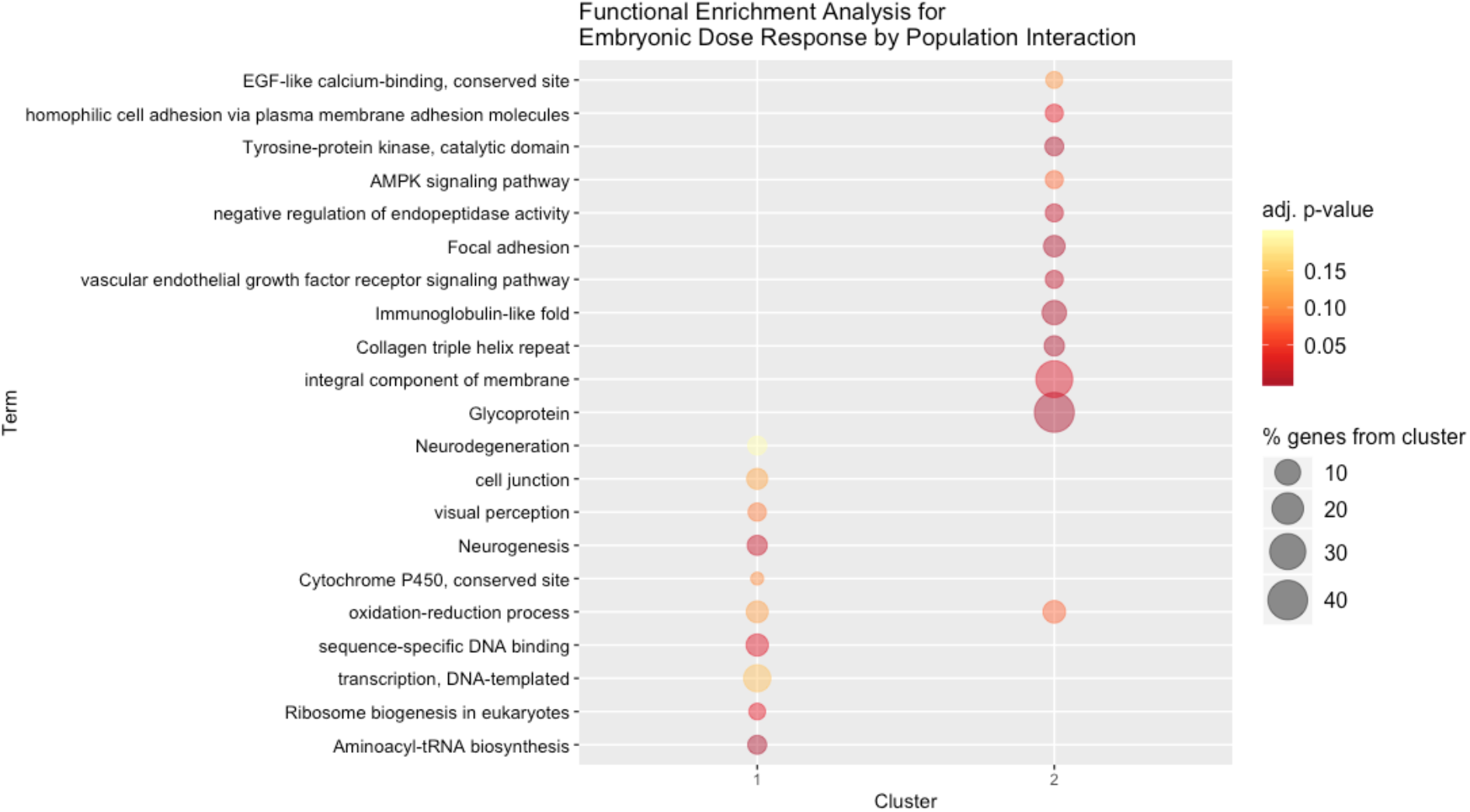
Gene Ontology (GO) enrichment analyses for the 512 genes that showed an oil exposure response that varied between populations (significant oil dose-by-population interaction, p<0.05). Cluster 1 represents functions enriched in the top cluster of DEGs in 5a (genes up-regulated in response to oil exposure in sensitive populations), and cluster 2 represents functions enriched in the bottom cluster of DEGs in 5a (genes down-regulated in response to oil exposure in sensitive populations).

**Figure S3.**
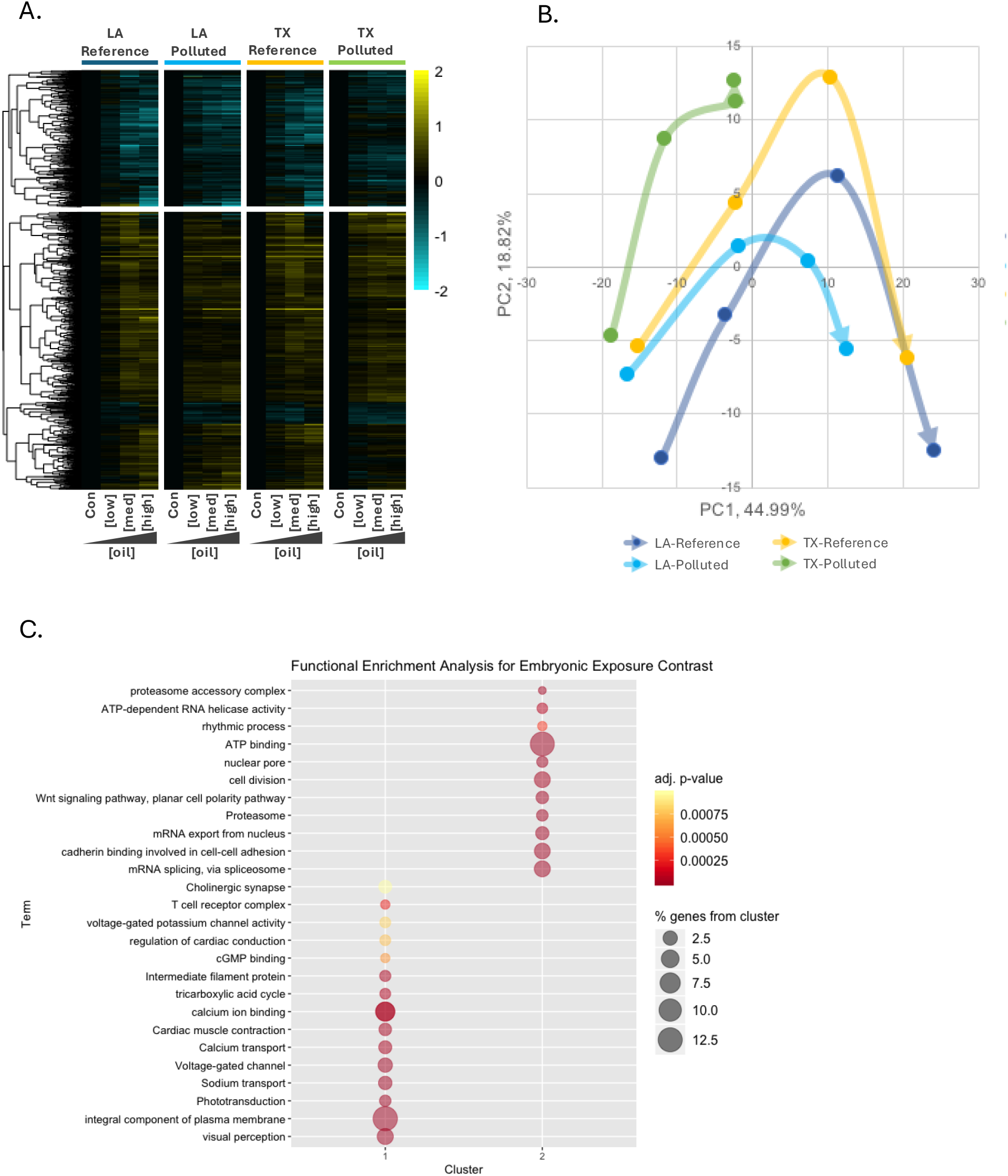
4,109 DEGs show a conserved transcriptional response to oil exposure in all four populations. We included genes with significant oil exposure response (FDR adj. p < 0.05) and excluded DEGs with a significant interaction effect of population by oil concentration (FDR adj. p < 0.1). A) Heatmap showing parallel (conserved) transcriptomic responses to oil between populations. The four panels, left to right, show oil concentration-responsive genes for the LA-Reference, LA-Polluted, TX-Reference, and Tx-Polluted populations, respectively. Within each population panel, increasing oil concentrations are organized starting from no-oil controls (Con) on the left to the highest concentration of oil on the right. Individual genes are the rows. Genes (rows) were hierarchically clustered (Pearson correlation). Higher and lower transcript abundance is indicated in yellow and blue, respectively. Color intensity relates to fold-increase or decrease of log2 expression (see color scale bar). B) The first two principal components for the oil exposure-responsive genes that were conserved in their response between populations. Arrows indicate the trajectory of gene expression change with increasing dose for each population. The base of each arrow represents the no-oil control condition for that population, where the arrow trajectory tracks gene expression change with increasing oil concentrations, such that the tip of arrow represents the highest oil concentration. C) The top and bottom clusters of co-expressed genes from the heatmap (cluster 1 and 2) were analyzed separately for GO enrichment analyses, which reveals biological pathways and functions involved in conserved responses to oil exposure.

